# The impact of the invasive *Kalanchoe* ×*houghtonii* on vegetated sea cliffs of the Mediterranean coasts with endemic *Limonium* species

**DOI:** 10.1101/2025.11.28.691185

**Authors:** Joan Pere Pascual-Díaz, Jordi López-Pujol, Neus Nualart, Sònia Garcia, Daniel Vitales

## Abstract

Invasive alien plant species pose a serious threat to biodiversity, especially in ecologically rich regions such as the Mediterranean Basin. Among the most affected habitats are coastal communities, which host many endemic taxa and provide valuable ecosystem services. One such invasive taxa is *Kalanchoe ×houghtonii*, an allegedly artificial hybrid. This plant, although having demonstrated strong invasive potential, still largely remains unrecognised as a taxon of concern in Mediterranean countries, at least in national official catalogues. In this study, we assess the impact of this hybrid on coastal communities, focusing on the European Union Habitat of Community Interest (HCI) 1240—“vegetated cliffs with endemic statices (*Limonium* spp.)”—at two sites along the southern coast of Catalonia (NE Iberian Peninsula). Between 2022 and 2025, we conducted field surveys to document the population size, growth stages, and spatial competition with native species. Additionally, we gathered 1,422 iNaturalist occurrences of *K. ×houghtonii* to map its distribution across the Mediterranean Basin, assess its presence within Natura 2000 protected sites and within protected areas, including the HCI 1240, and determine its potential spread using ensemble ecological niche modelling with bioclimatic variables. Our results show that *K. ×houghtonii* forms dense monospecific patches in the surveyed areas that compete for space with two native *Limonium* species in southern Catalonia. Moreover, we confirmed 713 naturalised occurrences in the Mediterranean area, 107 were located within Natura 2000 protected sites and 58 within the HCI 1240 included in protected sites. Ecological niche modelling indicates high climatic suitability across 93% of western and 59% of eastern Mediterranean Natura 2000 sites containing the HCI 1240. The findings of this study highlight the invasive potential of *K.* ×*houghtonii* and support its inclusion in national catalogues of invasive species across Mediterranean basin countries. The study calls for systematic monitoring of the spread and ecological impact of this hybrid species in coastal community habitats.

## INTRODUCTION

Invasive alien species (IAS) are animals, plants or micro-organisms introduced—whether accidentally or deliberately, but always as a result of human activities—into a natural environment where they are not naturally found, with significant impacts on local ecosystems (Pysek *et al*. 2020). Invasive species have become components of the floras and faunas across the world (van Kleunen *et al*. 2015) and are regarded as drivers of native species extinctions worldwide (Bellard *et al*. 2016), rising as the second cause of species gone completely extinct since 1500. In plants, invasive species have been associated with the extinction of up to 27% of species currently listed as extinct (Bellard *et al*. 2016). The importance of addressing IAS is reflected in one of the action-oriented global targets of the Kunming-Montreal Global Biodiversity Framework (target 6), which asks for reducing the rates of introduction and establishment of IAS by at least 50% by 2030 (Convention on Biological Diversity 2022).

Plant invasions are not evenly distributed across the world. The richness of alien floras depends on many factors generally related to human activities, such as trade volume, gross domestic product per capita, population density, habitat disturbance, and climate change (Essl *et al*. 2019). Europe, particularly its Mediterranean part, is amongst the world regions with the highest invasion potential (Bellard *et al*. 2016) and invasion threat (Early *et al*. 2016). The Mediterranean Basin, also a biodiversity hotspot (Mittermeier *et al*. 2011), has a history marked by the emergence of civilisations and trade commerce with other areas (since the Neolithic; Robb and Farr 2005) that, coupled with high population densities, have facilitated the introduction of new alien species. However, the rise of colonialism from the 16th century onwards, along with the development of long-distance communications in the 19th century, and the current globalisation, significantly increased the introduction of invasive species in Mediterranean-type ecosystems (Lambdon *et al*. 2008, MartínLForés 2017, Hulme 2021, Munné-Bosch 2023). Agriculture has always been one of the most important land and habitat-modifying activities, but nowadays the Mediterranean landscape is being dramatically affected by the growth of urban areas, the loss of rural environments, and the impact of tourism (Colaninno 2012). Some of the habitats most affected are coastal communities (Tordoni *et al*. 2020, Lami *et al*. 2021).

Although covering a relatively small area—due to its mainly linear nature—coastal habitats contain numerous endemic species and provide important ecosystem services (i.e., erosion regulation, biodiversity conservation, recreation) (Médail and Verlaque 1997, Blondel and Médail 2009). On the other hand, its often-intricate orography hinders the eradication of invasive alien plant species, which can easily colonise new sites from the highly anthropised surrounding areas. In this context, assessing the occurrence of invasive plant species in protected habitats and their potential impact on threatened native species is key to implementing effective conservation measures.

One of the most recently recorded neophytes in the Mediterranean Basin is *Kalanchoe* ×*houghtonii* (Crassulaceae) (Guillot-Ortiz *et al*. 2014). This taxon, of hybrid origin, was first reported by the horticulturist A.D. Houghton in the mid-1930s in California, from an artificial cross between *Kalanchoe daigremontiana* and *K. delagoensis*, two species native to Madagascar (Houghton 1935). This invasive hybrid taxon has been widely studied both from morphological (Shtein *et al*. 2021) and genomic (Pascual-Díaz *et al*. 2025) perspectives. Four different morphotypes (morphotypes A and B are purported to be artificial, while C and D may have a natural origin in Madagascar; Shtein *et al*. 2021) and three different cytotypes (2x, 3x, and 4x) have been reported, being tetraploid plants—the so-called morphotype A—the most successful invaders (Pascual-Díaz *et al*. 2025). While morphotype B is increasingly found at the global level, morphotypes C and D would still be confined to the island. This invasive capacity is enhanced by the spread of new clonal individuals through plantlets emerging from leaf margins (Shtein *et al*. 2021, Pascual-Díaz *et al*. 2025). While this reproductive trait is shared with its parental species, they exhibit lower colonisation capacity as compared to the hybrid (Guerra-Garcia *et al*. 2015, Herrera *et al*. 2018).

The presence of wild populations of *K.* ×*houghtonii* was first reported in Australia (1965) and the Bahamas (1970). Since then, this hybrid has shown a worldwide expansion and is now present in all continents except Antarctica (Herrando-Moraira *et al*. 2020). Despite its widespread distribution, *K.* ×*houghtonii* is officially listed as an invasive species only in the Australian state of Queensland (Biosecurity Act 2014, Australian Queensland Government) and in the US state of Florida (Florida Exotic Pest Plant Council, 2017). However, although *K.* ×*houghtonii* has not been recognised as an invasive species in official (i.e., under regulation) catalogues elsewhere, the Mediterranean Basin—particularly coastal habitats—is one of the most affected regions by its presence (Herrando-Moraira *et al*. 2020). The first confirmed wild record in the Mediterranean Basin is that of Alacant Province in eastern Spain in 1993, which was initially identified as *K. daigremontiana* (Guillot Ortiz *et al*. 2015) till recently, when we had the chance to access the herbarium sheet. Since then, this taxon is increasingly present in mostly scientific, non-official checklists of alien flora, such as those from Algeria (Sakhraoui and Thomson 2024) or Italy (Galasso *et al*. 2024). In Spain, the genus *Kalanchoe* is included in the checklist of ‘allochthonous species liable to compete with native wildlife, alter their purity, or disrupt ecological balances’ (Spanish Government 2025). However, *K.* ×*houghtonii* has not yet been officially included in any legally binding list of invasive species across the Mediterranean, which hinders the implementation of control and eradication measures by public administrations.

The absence of formal regulatory frameworks in the Mediterranean countries is probably due to its recent introduction, as compared to other regions such as Australia or North America (Herrando-Moraira *et al*. 2020). The scarcity of comprehensive data on the ecological impact of *K. ×houghtonii* in the Mediterranean region has further impeded its inclusion in official regulatory listings. Nonetheless, an increasing body of evidence suggests that *K. ×houghtonii* exhibits clear invasive potential in natural ecosystems of high conservation value, particularly in Mediterranean coastal habitats rich in endemic flora (e.g., in the Balearic Islands, Palerm *et al*. 2011; or in the south-eastern Iberian coast, Lahora *et al*. 2017). The presence of this hybrid taxon in the Iberian Peninsula has shown an exponential increase in recent years, reflecting a dynamic process of range expansion. Guillot Ortiz *et al*. (2014) documented 62 georeferenced occurrences distributed throughout the Iberian Peninsula and the Balearic Islands, highlighting its relatively limited but already established presence at that time. Only 16 years later Herrando *et al*. (2020) reported 177 occurrences for Catalonia, at the NE of the Iberian Peninsula. This trend continued in the following years, with the identification of 32 additional localities in the small area of the Costa Brava in NE Catalonia (Gómez-Bellver *et al*. 2025). These data suggest a rapid and ongoing expansion of the hybrid’s distribution range, particularly along the Mediterranean coast, and possibly facilitated by favourable climatic conditions, human-mediated dispersal, and ecological adaptability (Utjés *et al*. 2021).

The ability of *K.* ×*houghtonii* to outcompete native species (Gómez-Bellver *et al*. 2025) has also drawn the attention of local naturalists at two coastal sites in southern Catalonia, who observed that the hybrid taxon was invading populations of native statices (*Limonium* spp.), species that play an important role in the sustainability of coastal and salty inland habitats (Doğan *et al*. 2020). These two sites are located within the Habitat of Community Interest (HCI) 1240, “vegetated sea cliffs of the Mediterranean coasts with endemic statices (*Limonium* spp.)”— part of the Habitats Directive of the European Union (Council Directive 92/43/EEC). The vegetation of this habitat is mainly composed of *Crithmum maritimum*, accompanied by several species of *Limonium* with a generally narrow distribution. In Catalonia, the HCI 1240 covers 595.36 ha (Government of Catalonia 2019), accounting for 22% of this habitat in Spain, and can be subdivided into two main zones: the north zone (from Portbou to Blanes), where this habitat forms an almost continuous band along the Costa Brava, and the south zone (from Castelldefels to northern Ebro Delta), with a more discontinuous and less uniform distribution (Balaguer, Gómez-Pujol and Fornós 2009). To gain insight into the interactions between *K*. ×*houghtonii* and native species in this habitat, we conducted targeted fieldwork in the two previously mentioned southern Catalan coastal sites.

Building on this field-based evidence of local impact, we extended our analysis to assess the broader distribution and invasion risk of *K. ×houghtonii* across the Mediterranean Basin, focusing on the HCI 1240 in general rather than on specific species. The conservation values of this habitat are manifold, including the high number of endemic and threatened plant species inhabiting it, sometimes exclusive of this habitat (*e.g. Armeria ruscinonensis*, in NE Spain-SW France; *Euphorbia margalidiana* in the Balearic Islands; *Limonium quinii* in Rhodes Island of Greece; or *Seseli farrenyi* in NE Spain), or the fact that it offers suitable nesting sites for endemic and/or threatened bird species such as the Balearic Shearwater (*Puffinus mauretanicus*) (Balaguer, Gómez-Pujol and Fornós 2009). Yet, the conservation status of this habitat is considered poor in the EU, needing thus changes in management and policy (European Environmental Agency, 2013).

Using citizen science data and an ecological niche modelling we aimed to (i) map the current occurrences of this hybrid plant in the Mediterranean region, (ii) identify its presence within Natura 2000 protected areas (PAs), and also within PAs that include the HCI 1240, and (iii) model its potential geographic distribution under current climatic conditions, with particular attention to the HCI 1240. This integrative approach allows us to connect local ecological impacts with regional invasion dynamics, providing a stronger foundation for future monitoring, risk assessment, and recommendation of regulatory measures.

## MATERIALS AND METHODS

### Fieldwork in Mont-roig del Camp and Tarragona (Catalonia, Spain)

To study the invasion dynamics of *K.* ×*houghtonii* within the HCI 1240 in the southern coast of Catalonia, we conducted four fieldwork campaigns between 2022 and 2025 in Miami Platja (Mont-roig del Camp) and Fortí de la Reina (Tarragona) (Figure 1), where local naturalists observed impacts of the hybrid on populations of *Limonium* spp. (J. R. Mendo Escoda, pers. comm.). Within Mont-roig del Camp, we prospected two localities: the northern side of Platja Cristall (41.00°N, 0.94°E) and Cala del Solitari (41.01°N, 0.94°E). In Tarragona, we surveyed the surrounding area of Fortí de la Reina (41.11°N, 1.27°E).

**FIGURE 1.**
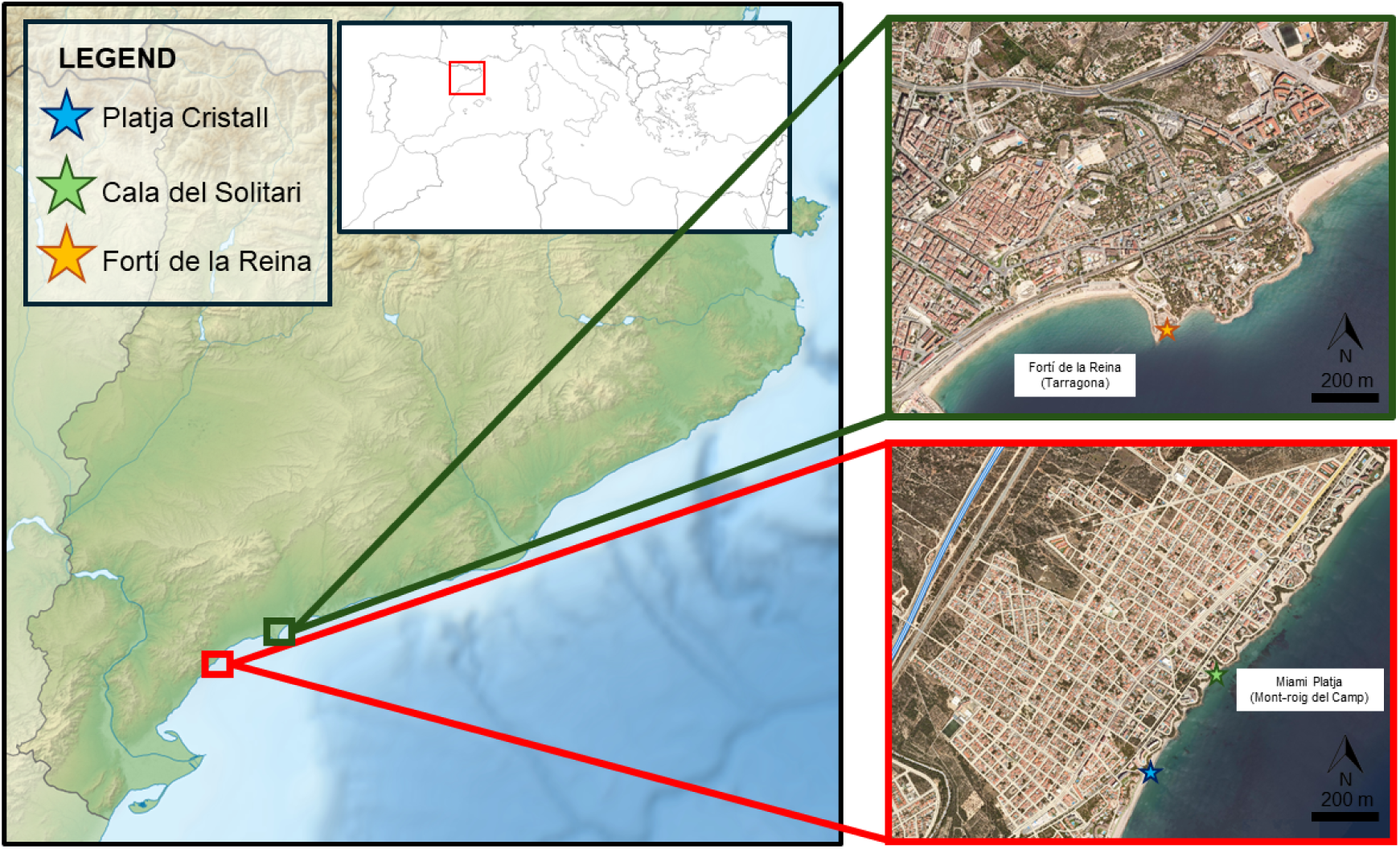
Detailed map of the three localities sampled along the coast of Miami Platja (Mont-roig del Camp) and Fortí de la Reina (Tarragona). In blue, the surveyed zone on the north side of the Platja Cristall; in green, the Cala del Solitari; in yellow, the prospected zone of the Fortí de la Reina. Credits of the maps: modified from Institut Cartogràfic i Geològic de Catalunya.

Fieldwork was conducted during the period of maximum vegetative activity (March–June) to facilitate species identification and the recognition of the phenological stages of *K.* ×*houghtonii*. We estimated the number of individuals (as visible shoots), measured the area occupied, and recorded the distribution of phenophases. Considering the difficulty of counting individuals of *K.* ×*houghtonii* because of its massive vegetative spread by propagules (implying that extremely dense monospecific patches of up to 1000–2000 individuals/m^2^ are often formed; Herrera *et al*. 2012; Tabares Mendoza 2016), the size of each population was inferred following an *ad hoc* approach. First, we counted in the field the number of cells of approximately 1 m^2^ where the species was present in the area. Then, we categorised the cells into two different types, defined based on former estimates (i.e., Herrera *et al*. 2012, Tabares Mendoza 2016) and our own in situ observations, as follows: (i) ‘low-density’ cells with *ca.* 250 individuals, and (ii) ‘high-density’ cells with *ca.* 1,000 individuals. For each cell, we estimated the proportions of the different phenophases, defined as follows: (i) young vegetative individuals < 5 cm height (considering 5 cm as the size at which they begin to produce propagules; Palerm *et al*. 2011), (ii) adult vegetative individuals > 5 cm height, and (iii) individuals with inflorescences (but where plantlets are also produced, usually larger and more successful than those developed on the leaves; Tabares Mendoza 2016). Additionally, we determined the presence of *Limonium* species for each locality with which *K.* ×*houghtonii* would potentially compete. Finally, for both Miami Platja and Fortí de la Reina localities, we also compiled a checklist of other invasive and potentially invasive alien species present in the area.

### Presence of *K. ×houghtonii* within Mediterranean protected areas and within PAs including the HCI 1240

The citizen science platform iNaturalist (https://www.inaturalist.org/) was employed to gather occurrence records along the Mediterranean Basin, due to several key advantages (López-Guillén *et al*. 2024): (i) each observation carries geographical precision, (ii) all observations are accompanied by images, which help verifying correct species identification and confirming that the specimens are wild rather than cultivated, (iii) it is, by far, the largest global platform for plant observations, and (iv) occurrence data are easy to download.

A total of 1,422 occurrences identified as *K.* ×*houghtonii* were extracted from iNaturalist in the Mediterranean Basin (defined as lat. SW: 30.22°, long. SW: -10.87°, lat. NE: 47.69°, long. NE: 38.17°), from the first reported record to 2 February 2025. Only those categorised as Research Grade (i.e., when more than two-thirds of identifiers agree on a taxon) were considered. Then, each occurrence was manually checked to exclude those that were in cultivation or escaped but not yet naturalised (e.g., for cases such as a few individuals occurring on roofs or floor cracks). Observations with low geographic precision (>300 m) and those where the hybrid morphotype was impossible to determine were also excluded from the study (Table S1). To ameliorate the sampling bias, the R package *spThin* was used to exclude too close occurrences (those that clustered within 50 m, as they probably belonged to the same population). After all the adjustments, a total number of 723 filtered observations along the Mediterranean Basin were used for downstream analyses.

To determine the presence of *K.* ×*houghtonii* in Mediterranean PAs, we first verified, using spatial data from the World Database on Protected Areas (WDPA, https://www.protectedplanet.net/), that none of the occurrences were located within a PA outside the Natura 2000 network. Then, we identified the filtered occurrences within Natura 2000 sites, based on spatial data from the European Environmental Agency. Finally, to assess the susceptibility of the HCI 1240 to *K.* ×*houghtonii* invasion, we determined which protected areas are associated with this particular coastal habitat. The most recently updated HCI 1240 map (https://biodiversity.europa.eu/habitats/ANNEX1_1240) has too low spatial resolution (∼10 km) to address this question accurately. Therefore, we overlapped *K.* ×*houghtonii* occurrences in the Mediterranean Basin with Natura 2000 sites that are known to contain the HCI 1240; such information was obtained from the Natura 2000 Viewer (https://natura2000.eea.europa.eu/). All spatial analyses were conducted in QGIS (v. 3.36.0, Maidenhead) and R (v. 4.1.2, R Core Team 2021).

### Modelling of the distribution area of *K. ×houghtonii* in the Mediterranean Basin

A suite of 19 bioclimatic variables was retrieved from the WorldClim 2.0 dataset (https://www.worldclim.org/) at 30 arc-seconds resolution, which were used for the modelling procedures. Pearson’s correlation coefficients were calculated among all bioclimatic variables to identify redundant predictors, only retaining those uncorrelated (*r□*<□|0.85|). For highly correlated variables, we calculated the median absolute correlation of each variable and retained those with the lowest median value. The final dataset of non-correlated bioclimatic variables included: mean diurnal range (bio2), temperature seasonality (bio4), max temperature of warmest month (bio5), mean temperature of coldest month (bio11), precipitation of wettest month (bio13), precipitation of driest month (bio14), precipitation seasonality (bio15), precipitation of warmest quarter (bio18), precipitation of coldest quarter (bio19). Afterwards, we extracted the values of the predictor variables for each occurrence point. To reduce the oversampling of the north-western Mediterranean (where the number of occurrences is much higher, probably reflecting higher iNaturalist activity and community engagement; Figure S1), we allowed only one occurrence per km^2^ (matching the spatial resolution of the environmental predictors, 30 arc-seconds ≈ 1 km at the equator), reaching a total number of 377 occurrences as input for the species distribution model. Two datasets of pseudoabsences with the same number of occurrences were placed randomly to calibrate the model. An ensemble of forecasts of species distribution models was then obtained, including projections from five statistical models: Generalised Linear Models (GLM), Generalised Additive Models (GAM), Multivariate Adaptive Regression Splines (MARS), Random Forests (RF), and Generalised Boosted Models (GBM). Models were calibrated using 70% randomly selected occurrence data, and accuracy was evaluated against the remaining 30% of the data, using the True Skill Statistic (TSS) (Allouche *et al*. 2006), and the area under the curve (ROC) (Swets *et al*. 1988). The analysis was replicated 10 times, thus providing a 10-fold internal cross-validation of the models. To summarise all projections into a meaningful integrated projection, we employed an ensemble strategy using committee averaging based on each model’s TSS values (greater than 0.35). Models and the ensemble forecasting procedure were performed within the R package *biomod2* (Thuiller *et al*. 2009).

To better understand the potential impact of *K. ×houghtonii* on Natura 2000 sites, as well as on the fraction of protected areas being composed of the HCI 1240, we used QGIS (v. 3.36.0, Maidenhead) to intersect the ensemble projections showing over 50, 70 and 95 percent of suitability for the presence of *K. ×houghtonii* with the GeoJSON file from the Natura 2000 Viewer (https://natura2000.eea.europa.eu/) containing all protected areas including the HCI 1240, only selecting those areas within the Mediterranean bioregion.

## RESULTS AND DISCUSSION

### Invasion dynamics of *Kalanchoe ×houghtonii* in the southern coast of Catalonia: Miami Platja (Mont-roig del Camp) and Fortí de la Reina (Tarragona)

The results from fieldwork along the southern littoral coasts of Catalonia are summarised in Table 1. At Miami Platja (Mont-roig del Camp), specifically in Cala del Solitari (Figure 2A), we observed the invasive hybrid covering approximately 400 m^2^, with an estimated population of 15,000 to 20,000 individuals, forming separate dense monospecific patches that partially dominate the soil cover. Most of the individuals were small plants, under 5 cm tall, but a considerable number were grown enough to start producing propagules. Larger individuals with inflorescences were in a lower proportion, being mainly restricted to the upper parts of the cliff, where they receive more sunlight, and propagules are released towards lower levels. Our demographic results, coupled with field observations, suggest that this population of *K.* ×*houghtonii* is currently threatening the viability of the native *Limonium virgatum* found at Cala del Solitari (Figure 2B, Figure 2D). The competition—primarily for space—caused by the intensive vegetative reproduction exhibited by *K.* ×*houghtonii* would be the main pressure for *Limonium* individuals in the site. The dense carpets formed by juvenile individuals of *K.* ×*houghtonii* significantly reduce the available space for other plants to grow. Over time, some of these *Kalanchoe* juveniles develop into mature individuals that, in turn, produce new propagules, further expanding the population. This reproductive strategy results in a population structure dominated by numerous juvenile individuals, interspersed with a sizable number of mature ones responsible of propagule production (Table 1). Given these dynamics and the potential further expansion of *K.* ×*houghtonii* throughout Cala del Solitari, not only could the persistence of *L. virgatum* be threatened in this locality, but the conservation status of the HCI 1240 could also be negatively affected. Populations of *K.* ×*houghtonii* can undergo significant expansion within just 5–6 years of initial establishment, a pattern observed in peri-urban areas of Barcelona (J. López-Pujol, pers. obs.) as well as on the Costa Brava in northern Catalonia, where some populations now occupy uninterrupted stretches of coastline spanning several hundred meters, with population sizes reaching several hundred thousand individuals (Gómez-Bellver *et al*. 2025). In addition to direct competition with native plant species, the hybrid might also affect the recipient ecosystems through its allelopathic effects, but also because of its ability to modify the carbon cycle in the soil, and alter both the direction of ecological succession and the composition and physiognomy of native-plant communities (as reported for *K.* ×*houghtonii* in Venezuela; Herrera *et al*. 2016).

**FIGURE 2.**
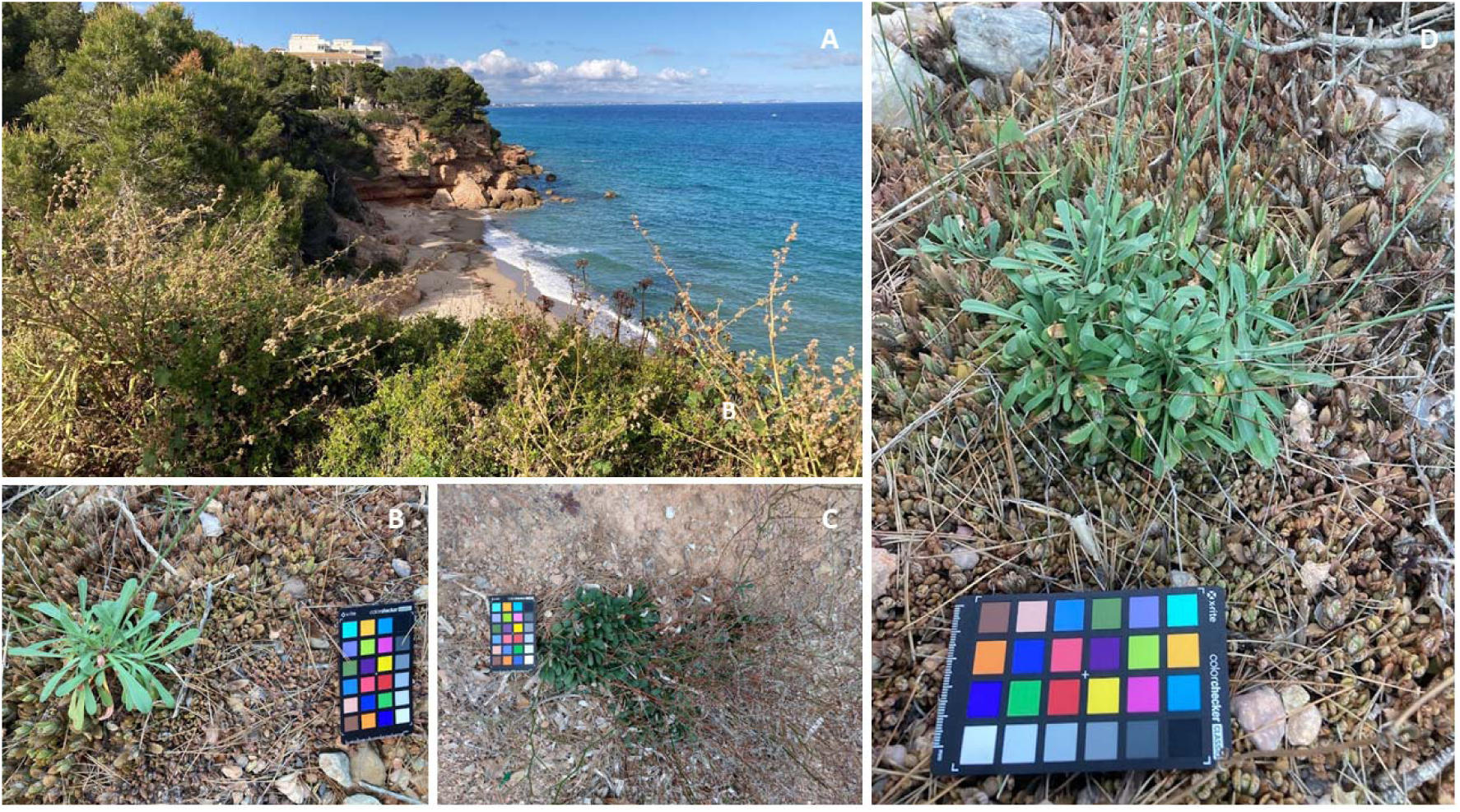
(A) Cliff ledge of Cala del Solitari (Mont-roig del Camp), where some *K.* ×*houghtonii* plants with inflorescences can be observed. (B) Individuals of *Limonium virgatum* surrounded by a dense patch of *K. ×houghtonii*. (C) Individuals of *Limonium gibertii*, a species included in the catalogue of threatened flora of Catalonia, from the locality of Platja Cristall. (D) Direct competition between *L. virgatum* and *K. ×houghtonii* in Cala del Solitari. Picture credits: Daniel Vitales.

**TABLE 1.** Summary of population size, area occupied, and phenological stages of *K. ×houghtonii* in Mont-roig del Camp and Tarragona.

|  | Mont-roig del Camp | Tarragona |
| --- | --- | --- |
| Number of individuals | 15,000–20,000 | 130,000–140,000 |
| Area occupied (m<sup>2</sup>) | 400 | 26,650 |
| Individuals < 5 cm | 65% | 60% |
| Individuals > 5 cm | 30% | 25% |
| Individuals with inflorescences | <5% | 15% |

In Platja Cristall—the second locality surveyed in Mont-roig del Camp—we detected the presence of *Limonium gibertii* (Figure 2C), a species endemic to the Balearic Islands and the eastern coast of the Iberian peninsula. It is listed as a protected species in Catalonia (officially classified as ‘vulnerable’; Diari Oficial de la Generalitat de Catalunya, CVE-DOGC-A-23325029-2023). Although this population is not currently threatened by *K. ×houghtonii*, we detected a large population of the hybrid nearby (less than 100 m apart; https://www.inaturalist.org/observations/203256812). Given its ease of spread, the *L. gibertii* population in Platja Cristall could be affected by *K. ×houghtonii* in the near future.

In Tarragona (capital), we surveyed the area surrounding the historical building of the Fortí de la Reina, a bastion built in the 18th century. There, we found a population of *K. ×houghtonii* roughly six times larger than the one in Mont-roig del Camp, covering an estimated area of 26,650 m^2^ and showing 130,000 to 140,000 individuals (Table 1). Again, the population predominantly consisted of individuals shorter than 5 cm, although a considerable number of individuals capable of producing propagules were also found. Compared to the population found in Cala del Solitari (Mont-roig del Camp), this locality had a higher proportion of individuals with inflorescences, potentially contributing to further extending the population by releasing larger and more successful plantlets (Tabares Mendoza 2016). The extension and abundance of *K. ×houghtonii* in this locality could also be related to its long-term occurrence. This population (Figure 3A) has been present for at least eleven years (according to historical imagery of Google Maps, see Figure S2; J. López-Pujol, pers. obs. in 2014), and may have existed for around 30 years (J.R. Mendo Escoda, pers. comm.). In the surveyed area, and similar to what we found in Cala del Solitari with *L. virgatum*, the dense patches of *K.* ×*houghtonii* are competing for the space with individuals of both *L. gibertii* and *L. virgatum* and, in many cases, the individuals of *Limonium* spp. were surrounded—and apparently displaced (some individuals interspersed with *K. ×houghtonii* were decaying)—by vegetative individuals of the hybrid (Figure 3B).

**FIGURE 3.**
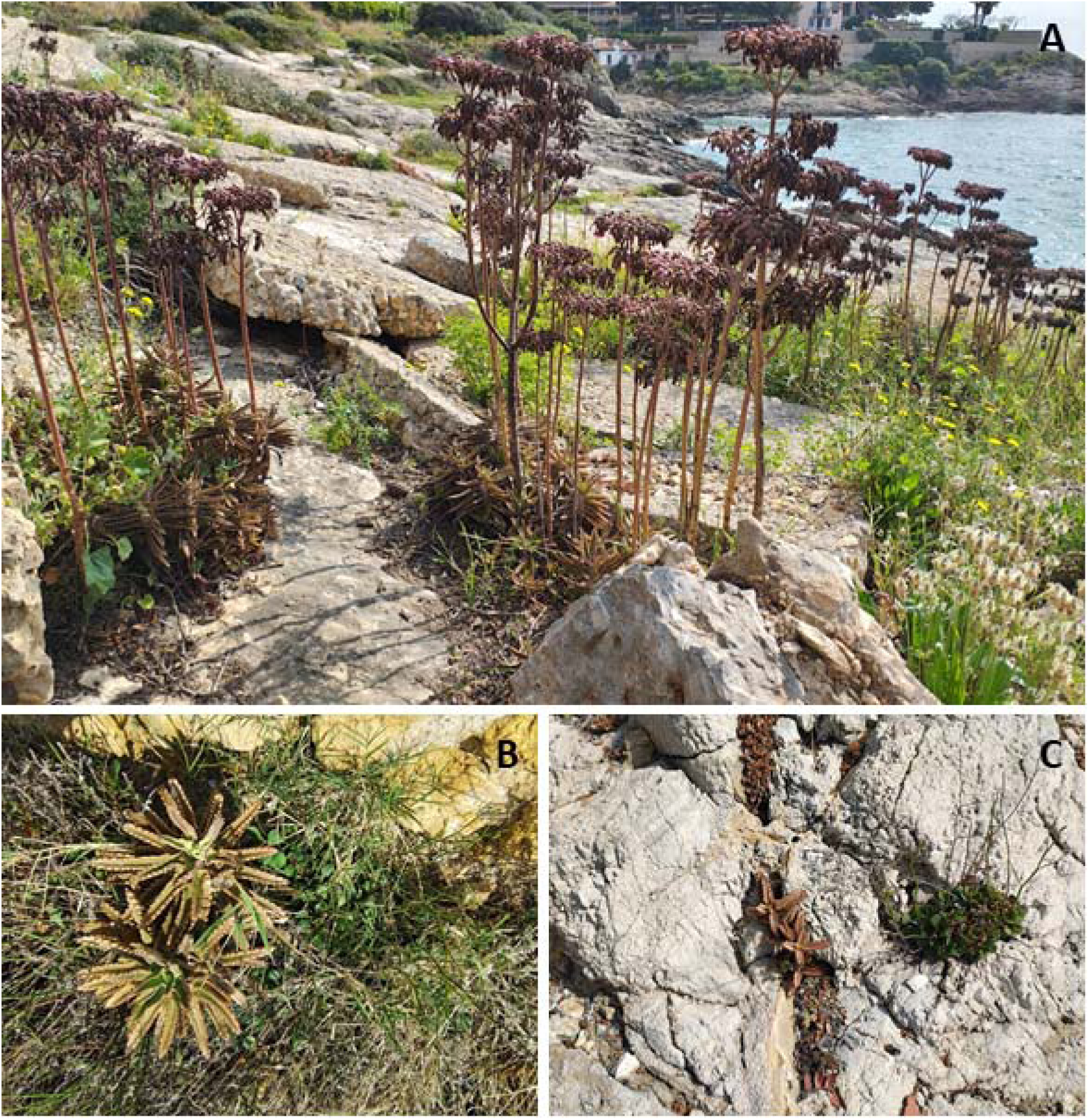
(A) Patches of individuals of *K.* ×*houghtonii* with inflorescences in the coastline surrounding the Fortí de la Reina (Tarragona). (B) Direct competition for the space of *K.* ×*houghtonii* with *Limonium virgatum* and (C) with *Limonium gibertii*. Picture credits: Jordi López-Pujol.

The invasive performance of *K. ×houghtonii* populations surveyed in the southern coast of Catalonia is consistent with those previously reported in other coastal areas of the Mediterranean Basin. In Eivissa island of the Balearic Archipelago (Palerm *et al*. 2011), *K. ×houghtonii* was initially detected in small, isolated patches in Ses Salines Natural Park, specifically in La Revista (a small rural settlement) and Es Cavallet (a beach area with nearby residential zones). However, over time, the plant began to spread aggressively, affecting an area over 9,000 m^2^ and threatening endemic and rare species such as *Limonium gibertii* and *Silene cambessedessi* (Palerm *et al*. 2011). In the Region of Murcia (Lahora *et al*. 2017), *K. ×houghtonii* was first confirmed naturalised at the cliffs of the Águilas Castle, competing for space and affecting native endangered species such as *Fumaria munbyi*, *Filago ramosissima* (not currently listed as protected in the Region of Murcia, although its inclusion in the regional catalogue has been recommended), and *Scrophularia arguta*. In the Valencian Country, in the Cap d’Or flora micro-reserve, a *Kalanchoe* species—most likely *K. ×houghtonii*, the only *Kalanchoe* reported in the area by iNaturalist and by our personal observations—has been largely recognised as a direct ecological threat to *Silene hifacensis* (Generalitat Valenciana, 2008), a flagship species and one of the most endangered endemic plants from Spain (Soler 2019).

These cases of invasion by *K. ×houghtonii* in the Iberian Peninsula and Balearic Islands share key ecological features: the invaded habitats are typically coastal, rocky environments—often cliffs or disturbed outcrops—with sparse native vegetation and usually high sun exposure, where *K. ×houghtonii* appears physiologically well adapted (Herrando-Moraira *et al*. 2020) and outcompetes native flora. Additionally, these sites are adjacent to urbanised or touristic areas, where propagule pressure is likely elevated due to gardening escape. This is further supported by the presence of many other non-native ornamental species co-occurring with *K. ×houghtonii* in the surveyed localities of Mont-roig del Camp and Tarragona (Table S2), indicating that the colonisation of coastal habitats is facilitated by nearby public and private gardens acting as propagule sources. In the two studied localities, as also occurs in other areas of the HCI 1240 in the Mediterranean Basin (Misuri *et al*. 2024; Gómez-Bellver *et al*. 2025), there are very rich assemblages of alien plants (up to 21 taxa in Tarragona; Table S2). Some of these alien plants are also competing and probably displacing the native *Limonium* species (Figure S3). Besides structural changes to communities, they also produce visual impacts that are also evident as some of the alien species are of big size (e.g., *Agave salmiana* subsp. *ferox*, *Opuntia stricta*), or occur in large quantities, forming eye-catching large carpets (e.g., *Carpobrotus* spp.). The massive entrance of alien plant species in the HCI 1240 may also produce a generalised native biodiversity loss, making these communities less resistant to further invasions (Tordoni *et al*. 2019).

### Distribution of *K.* ×*houghtonii* in the Mediterranean Basin

In the Mediterranean Basin, a total of 723 naturalised occurrences have been recorded through citizen science via iNaturalist, with Spain being the country holding the largest number of occurrences of the invader *K.* ×*houghtonii* (77.00%), followed by Italy (10.65%) and Portugal (4.84%). With a lower frequency, observations of the hybrid in the wild have also been detected in Algeria, Croatia, France, Greece, Jordan, Malta, Montenegro, Morocco, and Turkey. All observations of the invasive hybrid included in our analyses corresponded to morphotype A, which, according to previous studies, is the most invasive morphotype of *Kalanchoe ×houghtonii* (Shtein et al. 2021, Pascual-Díaz et al. 2025). However, the presence of the morphotype B is not discarded in the Mediterranean Basin, since it was detected close to the natural reserve of the Cap de Creus (Spain) (Pascual-Díaz *et al*. 2025), and in Ostia (Italy; https://www.inaturalist.org/observations/162486856), although in these two localities the plant is not yet naturalised.

There is a major west–east decreasing trend in the number of wild observations of *K. ×houghtonii* in the Mediterranean Basin (Figure 4), a pattern that could be explained by different factors. Firstly, the earliest documented occurrence of *K. ×houghtonii* in the Western Mediterranean region dates back to 1993 (Guillot-Ortiz *et al*. 2015, Pascual-Díaz *et al*. 2025), whereas the first confirmed record in the Eastern Mediterranean—specifically in Greece—was much more recent, in 2016, based on a citizen science observation (https://www.inaturalist.org/observations/7405228). Therefore, assuming that the year of first observation could reflect the actual timing of arrival, the hybrid taxon may have had much more time to expand in the western than in the eastern Mediterranean. Secondly, the highest number of observations in iNaturalist from western Mediterranean countries (Figure S1) could reflect a more intense iNaturalist participation rather than a real higher presence of the species; in other words, the occurrence of a given species could be underestimated in countries in which people use iNaturalist much less. Thirdly, it should be noted that citizen scientists can be biased towards charismatic, easily recognisable taxa (Petersen *et al*. 2021, and references therein); thus, species that are already known to the public (e.g., through gardening or media) are more easily recognised and therefore more often reported; in Spain, *Kalanchoe ×houghtonii* is a very popular plant widely sold in gardening stores (and is also very common as a plant exchanged among friends or acquaintances) because of its supposed anti-cancer properties. Fourthly, a better conservation status of the HCI 1240 in the eastern Mediterranean compared to the western Mediterranean Basin (European Environmental Agency, 2013) has been reported. Indeed, the distribution of *K. ×houghtonii* in the Mediterranean Basin is consistent with results from Cao Pinna *et al*. (2024), which in the current climatic conditions, France, Italy, Portugal, and Spain are the most affected areas in the European Mediterranean coast by the invasion of alien naturalised flora; therefore, the presence of the hybrid would also be consistent with the conservation status of the area which colonizes. Fifth, and last, bioclimatic conditions in the eastern Mediterranean may be less favourable for the introduction and establishment of *K. ×houghtonii* than in the western Mediterranean (discussed further in the next section).

**FIGURE 4.**
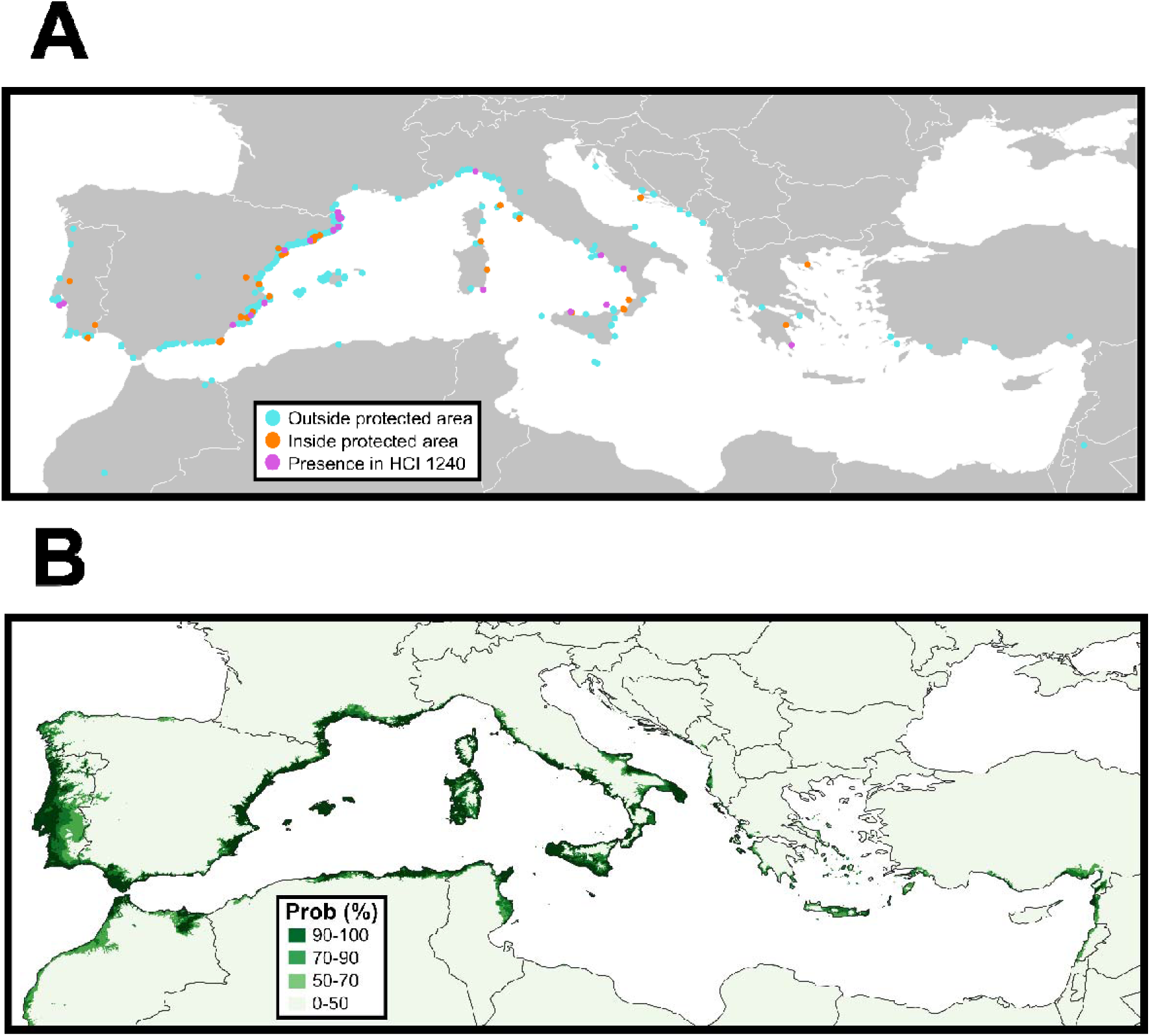
(A) Map of *K.* ×*houghtonii* occurrences in the Mediterranean Basin. In blue and orange, occurrences outside and inside the Natura 2000 protected areas, respectively. In purple, occurrences inside Natura 2000 protected areas and in the habitat of community interest ‘Mediterranean cliffs with endemic *Limonium* spp.’ (HCI 1240) (B) Projections of the ensembled potential distribution models of *K.* ×*houghtonii* in the Mediterranean Basin. Colours denote probabilities from 0% (soft green) to 100% (dark green).

Most of the factors that explain the west–east gradient in the number of *K. ×houghtonii* occurrences also account for the marked differences between the northern and southern parts of the basin: the plant arrived much later in the region (the first confirmed record dates to 2008 in Tunisia; Herrando-Moraira et al. 2020), iNaturalist is far less widely used, the species is little known to the general public, and the bioclimatically suitable area is very limited—restricted mainly to the coastal zones of Morocco, eastern Algeria, and northern Tunisia, while its presence in Libya and Egypt appears highly unlikely. An additional factor is the comparatively lower gross domestic product (GDP) of the southern basin countries, as economic development is a widely recognised key driver of biological invasions (Hulme 2009; Pyšek et al. 2010).

From all observations of *K. ×houghtonii* in the Mediterranean, 107 (14.80%) have been reported inside protected areas. More than half of the occurrences inside protected areas are also found within areas containing the HCI 1240 (i.e., a total number of 58 occurrences representing 8.02% of the whole dataset; Figure 4A). This strongly suggests that this habitat may be particularly sensitive to invasion by *K. ×houghtonii*. Rocky habitats are among the commonest in which the hybrid occurs around the world (Herrando-Moraira et al., 2020), likely mirroring those of the parental species in Madagascar, where they grow on granite, sandstone or limestone outcrops, or coastal and inland unconsolidated sands (Boiteau and Allorge-Boiteau 1995, Descoings 2003, Witt and Rajaonarison 2004), often in direct sunlight (Hannan-Jones and Playford 2002) and having adaptations to high irradiation (Kluge and Brulfert 2000). These conditions (rocky substrates and full sun exposition) are generally fully met in coastal cliffs. In addition, *K.* × *houghtonii* significantly increases plantlet production under experimental high light conditions (Zhang *et al*. 2024), which could enhance its colonization capabilities.

Most of these observations of *K.* ×*houghtonii* dwelling in the HCI 1240 inside PAs are concentrated in the western Mediterranean, with just a single observation in the eastern part of the basin (Greece) (Figure 4A). This geographic pattern matches the conservation status of the HCI 1240 in the Mediterranean Basin, being unfavourable in Spain, France, and Italy, but favourable in Croatia and Greece (https://nature-art17.eionet.europa.eu/article17/habitat/summary/?period=5&subject=1240). This can be linked to the hypothesis of biotic resistance to non-native species in speciesLrich communities (Rejmánek 1996), which would be those in a better state of conservation. Indeed, the relatively large number of field studies documenting the invasion of *K.* × *houghtonii* in western Mediterranean coastal areas (*e.g.* Palerm *et al*. 2011; Lahora *et al*. 2017; Gómez-Bellver *et al*. 2025; this study) could be interpreted as indirect evidence of the poorer conservation status of HCI 1240 in this part of the basin.

### Projection of the potential habitat of *Kalanchoe* ×*houghtonii* in the Mediterranean Basin

To investigate the potential for future spread and identify areas at risk—especially within or near PAs—we conducted species distribution modelling to estimate the climatically suitable range of *K. ×houghtonii* across the Mediterranean Basin. The projection of the potential distribution of *K.* ×*houghtonii* indicates that the western Mediterranean—particularly Spain and Italy—is highly suitable for the nothospecies under current climatic conditions, especially in coastal and peri-urban areas. Although recorded occurrences are currently much lower in the eastern Mediterranean (FigureL4A), ensemble species distribution models suggest that many coastal areas in this region also present favourable climatic conditions for its establishment (Figure 4B). Given the still low number of occurrences and the climatic suitability of the hybrid, we consider coastal areas of the eastern Mediterranean Basin as a “red alert” zone—*i.e.*, an area not yet widely colonised but highly vulnerable to future invasions (Herrando-Moraira *et al*. 2020).

Across the European Mediterranean bioregion, there are 433 Natura 2000 PAs which include the HCI 1240. Of these, 73.21% are located in the western Mediterranean, while 26.79% are in the eastern Mediterranean. According to our distribution model (at a suitability threshold of 50%), 93.34% of western Natura 2000 PAs containing the HCI 1240 are suitable for colonisation by *K.* ×*houghtonii*. In the eastern Mediterranean, 59.36% of such areas are suitable. This lower percentage in the east is partially due to the Natura 2000 PAs of Slovenia—four of which harbour the HCI 1240, none of them being inferred as climatically suitable for the hybrid. In contrast, Croatia, Greece, and Cyprus show suitability rates of 68.75%, 97.26% and 71.43%, respectively, across all their Natura 2000 PAs containing the HCI 1240.

When the suitability threshold is increased to 70%, the potential presence of *K.* ×*houghtonii* in Natura 2000 PAs in the western Mediterranean remains unchanged, while it slightly decreases in the eastern Mediterranean (42.83%). Finally, at a stricter suitability threshold of 90%, the western Mediterranean remains highly suitable for *K.* ×*houghtonii* (92.28%), whereas suitability in the eastern Mediterranean decreases substantially (26.20%). This reduction is mainly explained by the absence of suitable climatic conditions in the PAs harbouring the HCI 1240 in Cyprus and Slovenia, as well as by a decrease of climatically suitable Natura 2000 PAs in Croatia and Greece (50.00% and 54.79%, respectively) under the strict 90% threshold (Table S4).

Our results highlight the need to develop region-specific strategies to manage *K.* ×*houghtonii* across the Mediterranean. In the western Mediterranean, where this invasive species is already widespread and affecting native endangered flora, management should focus on mitigating the expansion of *K.* ×*houghtonii* and locally eradicating it within natural PA. In the eastern Mediterranean, where the hybrid taxon is currently less established according to available occurrence data, prevention efforts are essential to limit its spread. Our potential distribution models indicate that even under very strict suitability thresholds (Table S4), half of the PAs harbouring the HCI 1240 in Greece and Croatia (countries that manage more than 90% of the Natura 2000 sites containing the HCI 1240 in the eastern Mediterranean) could still be affected by *K.* ×*houghtonii*. Meanwhile, on the south-western part of the Mediterranean Basin, scarcity of records associated with limited botanical documentation and reduced citizen science activity, combined with the high habitat suitability in coastal areas (Figure 4B), indicates that this area should also be considered a “red alert” zone.

## Conclusions

Despite growing evidence of the current and potential impact of *K.* ×*houghtonii* on Mediterranean coastal habitats, this taxon is not yet included in the invasive species regulations of any country in this region. In Spain, the most recent catalogue on invasive species (Royal Decrees 630/2013 and 216/2019, Orders TED/1126/2020 and TED/339/2023) includes a total number of 199 invasive taxa (114 animals, 69 plants, 15 algae, and one fungus). However, *K.* ×*houghtonii* and their parental species, also known to exhibit invasive behaviour, are not included in this catalogue. Considering the ecological impacts on native species and habitats reported here and in previous studies (e.g., Palerm *et al*. 2011, Lahora *et al*. 2017, Gómez-Bellver *et al*. 2025), *K. ×houghtonii* meets the criteria for classification as an invasive species under Spanish legislation. Its formal inclusion in the Spanish Catalogue of Invasive Species would facilitate coordinated monitoring, control, support the implementation of eradication measures, and help mitigate further ecological degradation in vulnerable habitats. In other western Mediterranean countries such as Portugal, France or Italy, although no data on the impact of *K. ×houghtonii* on the local flora are yet available, the hybrid’s widespread occurrence and large potential distribution underscore the need to consider similar legal and management strategies.

The findings of this study also suggest that *K.* ×*houghtonii* should be considered for inclusion in checklists and catalogues of invasive species across eastern Mediterranean Basin countries. In Greece, the most recent list of alien plants is the *Atlas of Alien Plants of Greece* (http://www.alienplants.gr), which currently lists 456 alien species but does not include any *Kalanchoe* taxa. A similar situation is observed in Turkey, where only two *Kalanchoe* taxa (*K. delagoensis* and *K. blossfeldiana*) are included among the country’s 340 alien plants (Uludağ *et al*. 2017). In Croatia, the national catalogue of alien species (https://invazivnevrste.haop.hr/) includes 3,171 species, among them *K.* ×*houghtonii* and other species of the genus. However, the hybrid is not listed in the national blacklist of invasive species (OG 15/2018, 14/2019 from the Croatian Government), so its cultivation, commercialisation, or introduction into nature is not regulated. Based on current evidence—showing that this hybrid taxon is already naturalised in protected natural areas—we advocate for systematic monitoring of the spread and ecological impact of *K.* ×*houghtonii* across the eastern Mediterranean area.

## ACKNOWLEDGEMENTS

We thank Joan Ramón Mendo Escoda and Víctor Álvarez López (GETE–Ecologistes en Acció) for the early detection of *K. ×houghtonii* populations in Cala del Solitari and Fortí de la Reina, and for their assistance in tracking them in these areas. We are also grateful to Llorenç Sáez for his help in identifying the two *Limonium* species, and to Neus Ibáñez for her valuable support during fieldwork. This work was supported by the Catalan Government (grant 2021SGR00315), the Spanish Research Agency (grants PID2020-119163GB-I00, CNS2023-143604, and PRE2021-097873 to J.P.P.-D.), and the European Union through the nature conservation LIFE program (grant LIFE20 NAT/ES/001223).

## AUTHORS CONTRIBUTIONS

D.V. and J.P.P.-D conceived the study. J.P.P.-D, J.L.-P, N.N., S.G., and D.V. conducted the fieldwork. J.P.P.-D, S.G., and D.V. analysed the data. J.P.P.-D and D.V. wrote the first version of the manuscript. All authors edited and approved the final version of the manuscript.

**TABLE S1.** List of all verified iNaturalist occurrences, including their corresponding link and associated information.

**TABLE S2.** List of surveyed invasive species in Mont-roig del Camp and Fortí de la Reina.

---

**FIGURE S1.**
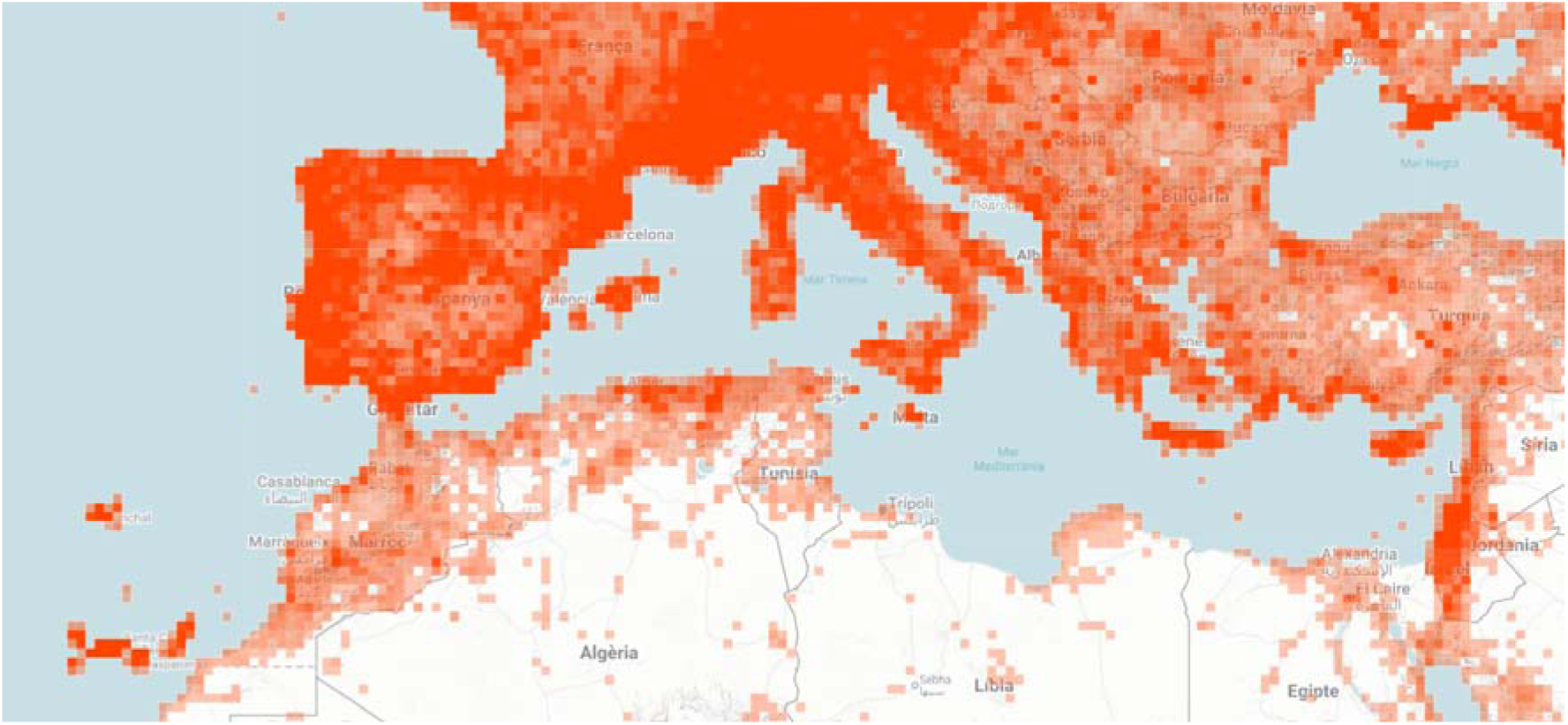
Screenshot (August 5, 2025) of the density of iNaturalist all plant observations (i.e., using the category ‘Plantae’ for search filter) in the Mediterranean Basin (source: https://www.inaturalist.org/observations?subview=map&taxon_id=47126). As iNaturalist does not offer public information on the geographic location of observers, we can take countries with the most observations as a good proxy for active participation as observers in this platform.

**FIGURE S2.**
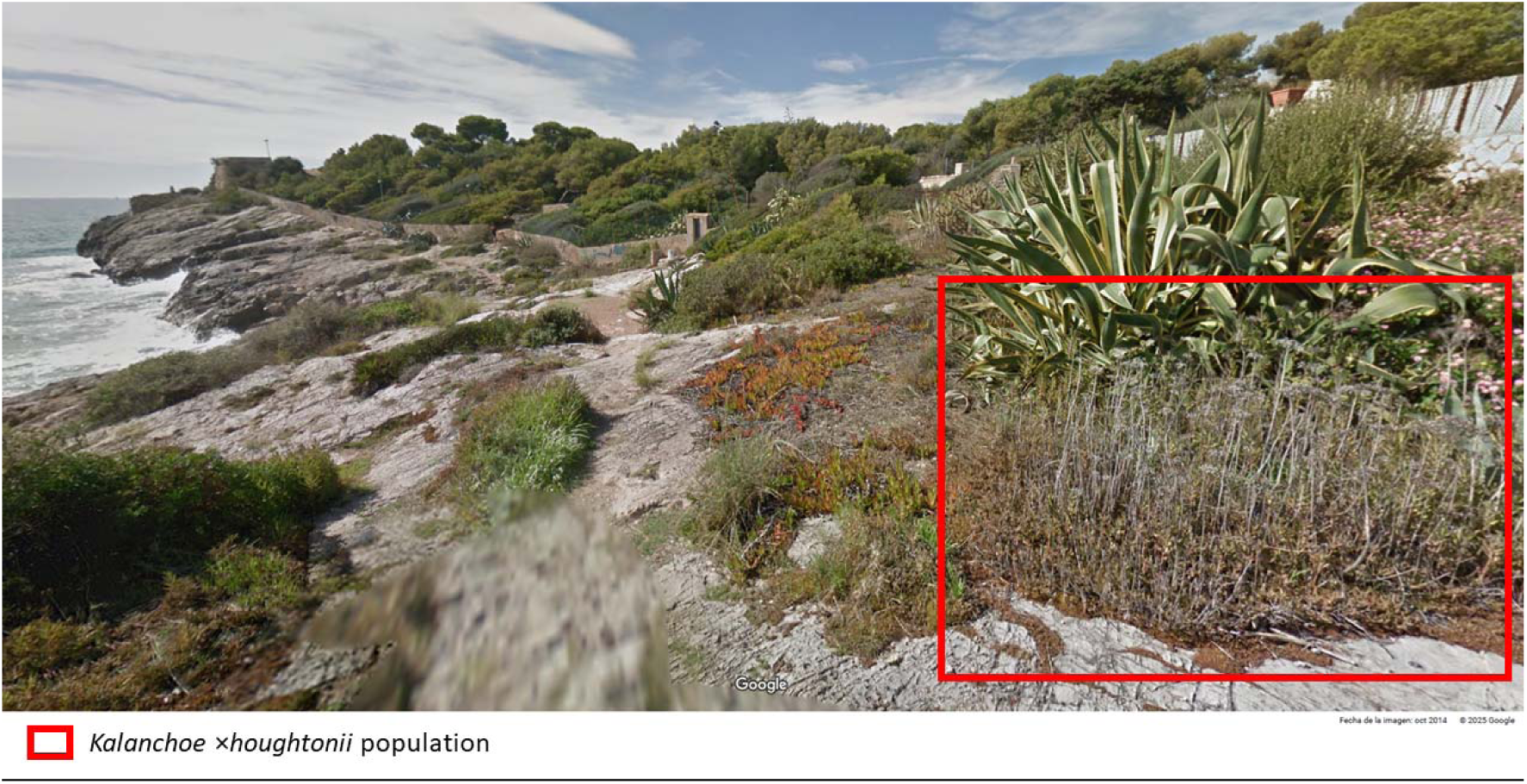
Picture extracted from Google Street View from October 2014 in the Fortí de la Reina (Tarragona, Catalonia, Spain) locality, with a population of *Kalanchoe ×houghtonii* established in the area.

**FIGURE S3.**
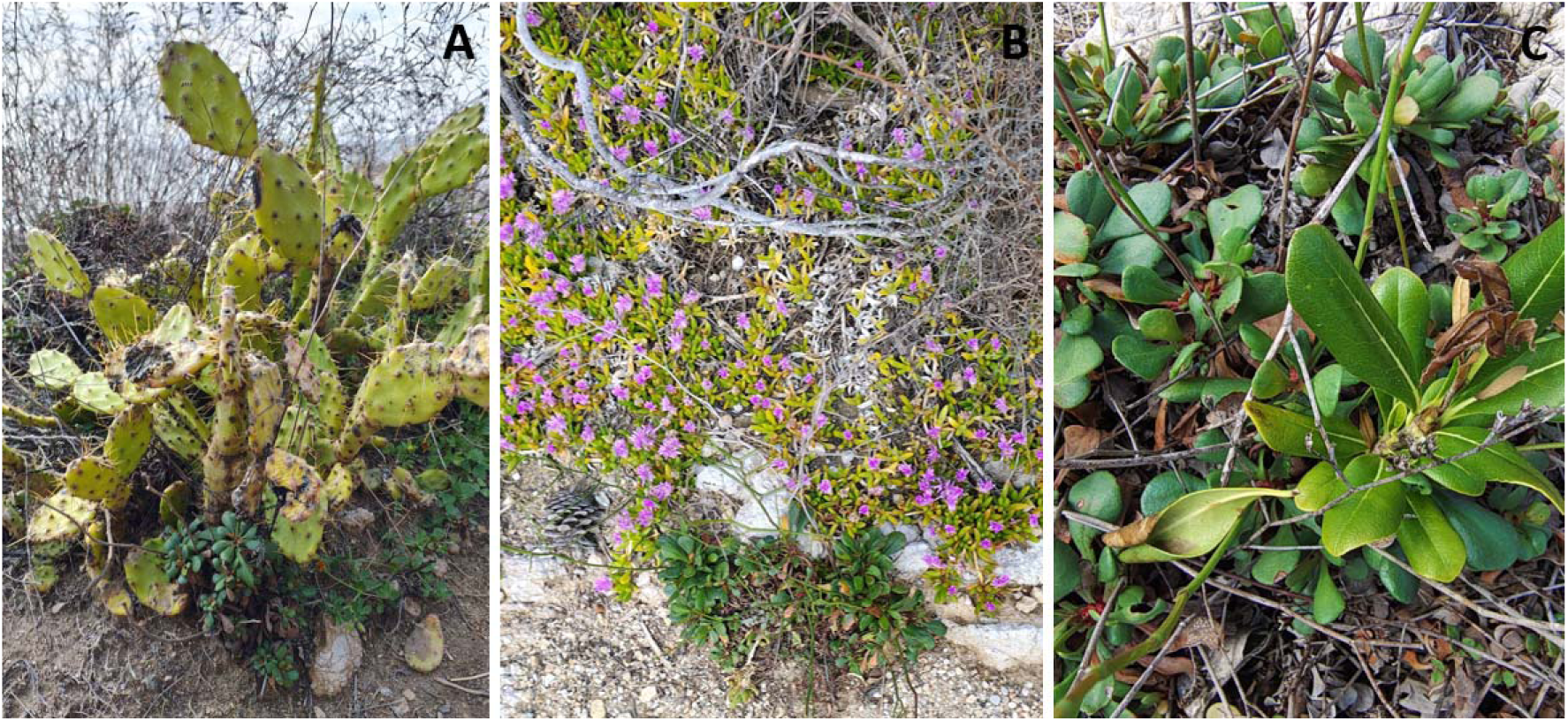
Pictures of (A) *Opuntia stricta*, (B) *Disphyma crassifolium*, and (C) *Pittosporum tobira* competing for space with *Limonium virgatum* (indicated with a white arrow) populations in Fortí de la Reina.

